# Anillin/Mid1p interacts with the ECSRT-associated protein Vps4p and mitotic kinases to regulate cytokinesis in fission yeast

**DOI:** 10.1101/2019.12.13.875211

**Authors:** Imane M. Rezig, Gwyn W. Gould, Christopher J. McInerny

## Abstract

Cytokinesis is the final stage of the cell cycle which separates cellular constituents to produce two daughter cells. Using *Schizosaccharomyces pombe* we have investigated the role of various classes of proteins involved in this process. Central to these is anillin/Mid1p which forms a ring-like structure at the cell equator that predicts the site of cell separation through septation in fission yeast. Here we demonstrate a direct physical interaction between Mid1p and the endosomal sorting complex required for transport (ESCRT)-associated protein Vps4p. The interaction is essential for cell viability, and Vps4p is required for the correct cellular localization of Mid1p. Furthermore, we show that Mid1p is phosphorylated by the aurora kinase Aurora A, that the interaction of *mid1* and *ark1* genes is essential for cell viability, and that Ark1p is also required for the correct cellular localization of Mid1p. We mapped the sites of phosphorylation of Mid1p by Aurora A and the polo kinase Plk1 and assessed their importance by mutational analysis. Mutational analysis revealed S332, S523 and S531 to be required for Mid1p function and its interaction with Vps4p, Ark1p and Plo1p. Combined our data suggest a physical interaction between Mip1p and Vps4p important for cytokinesis, and identify phosphorylation of Mid1p by aurora and polo kinases as being significant for this process.

**Author summary:** Replication is a property of all living cells, with cell separation, so-called cytokinesis, the final step in the process. A large number of proteins have been identified that are required for cytokinesis, but in many cases it is not understand how they interact and regulate each other. In this research we have analysed two classes of proteins founds in all eukaryotic cells with central roles in cytokinesis: the endosomal sorting complex required for transport (ESCRT) proteins and the anillin protein Mid1p. We identify a direct physical interaction between the ESCRT protein Vps4 and anillin/Mid1p, and explore how it regulates cytokinesis. Midp1 activity is shown to controlled by the protein kinase Ark1p by direct phosphorylation, and this phosphorylation is important for Mid1p function. These observations identify new ways in which ESCRT and anillin/Mid1p control cell separation.

## Introduction

Much is understood about the control and regulation of the cell division cycle, with many critical cell cycle mechanisms identified being evolutionarily conserved, present across the eukaryotic kingdom. One cell cycle stage that has been increasingly studied is cytokinesis where DNA, organelles and cell constituents are partitioned and allocated to two daughter cells during their physical separation. The fission yeast *Schizosaccharomyces pombe* has proven to be an excellent model organism to study the eukaryotic cell cycle [1]. It is especially useful for studying cytokinesis and cell division as, explicit in its name, it divides by medial fission involving a contractile actomyosin ring leading to the process of cell separation, which is similar to these processes in mammalian cells [2].

A number of proteins have been identified that regulate cytokinesis in fission yeast, including those in a signal transduction pathway named the Septation Initiation Network (SIN) [3]. The SIN proteins form a pathway that facilitates contractile ring constriction and promotes the formation of a medial cell wall-like structure between the two daughter cells, called the septum. Furthermore, SIN components associate with the spindle pole bodies and link mitotic exit with cytokinesis [4]. Sid2p is one regulator of the SIN-cascade of signaling proteins [5]. It terminates the signaling cascade leading to the transition from the spindle pole bodies to the cell division site promoting the onset of cytokinesis [6]. A recent study identified anillin/Mid1p as a substrate for Sid2p and indicated that the phosphorylation of Mid1p facilitates its removal from the cell cortex during the actomyosin contractile ring constriction [7].

Mid1p forms a ring-like structure at the cell equator that predicts the site of cell separation through the formation of a septum in fission yeast [8,9,10]. In cells containing a chromosomal deletion of the *mid1* gene (*mid1Δ*), a misshaped contractile ring is assembled during anaphase when the SIN pathway becomes active; such observations confirm the important role of Mid1p in directing contractile ring assembly to its correct location [11,12,13].

Many other groups of proteins have a role in septum formation in *S. pombe*. Amongst these the various classes of ESCRT proteins, including the ESCRT-III regulator Vps4p, were found to be required for septation, suggesting that they have a role in cytokinesis in fission yeast [14, 15]. Additional experiments suggested that the ESCRT proteins interacted with established cell cycle regulators including the polo kinase Plo1p, the aurora kinase Ark1p and the CDC14 phosphatase, Clp1p to control these processes [14].

As a way to further understand the regulation of ESCRT proteins during cytokinesis in fission yeast, we sought to identify other proteins that interact with anillin/Mid1p. Here we describe a direct physical interaction between Vps4p and the anillin Mid1p which is essential for cell viability and for the correct placement of Mid1p. We further show an interaction between Mid1p and the aurora kinase Ark1p which is essential for cell viability and identify phospho-acceptor sites in Mid1p and study their role using mutagenesis. Collectively, our observations reveal novel mechanisms by which cytokinesis is regulated by different classes of proteins acting through Mid1p.

## Results

### Genetic and physical interactions between Mid1p and Vps4p in fission yeast

Previously, we identified and characterised the requirement and role of ESCRT proteins for cytokinesis in fission yeast [15]. These experiments also revealed the interaction of ESCRT proteins with three cell cycle regulators, the Polo-like kinase Plo1p, the aurora kinase Ark1p and the CDC14 phosphatase Clp1p. These observations offered a framework by which ESCRT proteins are regulated to control cytokinesis. To further explore ways in which the ESCRT proteins might integrate with other cell cycle regulators to control cytokinesis, we examined their interaction with the anillin Mid1p. Mid1p has a central structural role in cytokinesis, forming equatorial nodes to create an annular shaped structure that determines the position of the division plane [16, 13].

We initiated this by searching for genetic interactions between the *mid1* gene and genes encoding ESCRTs and ESCRT-associated proteins. Double mutant fission yeast strains were created containing a chromosomal deletion of *mid1* (*mid1D*) and individual chromosomal deletions of ESCRT genes from Classes E-0 to E-III and *vps4* and searching for synthetic phenotypes. *mid1*Δ was combined with *sst4*Δ (E-0), *sst6*Δ and *vps28*Δ (E-I), *vps36*Δ and *vps25*Δ (EII), *vps20*Δ, *vps32*Δ and *vps2*Δ (E III), and *vps4*Δ [17, 15]. In each case double mutants were created which were viable, with no apparent synthetic phenotypes (data not shown). The exception was *mid1Δ vps4*Δ which instead failed to form viable colonies and was synthetically lethal (Fig 1A).

**Fig 1.**
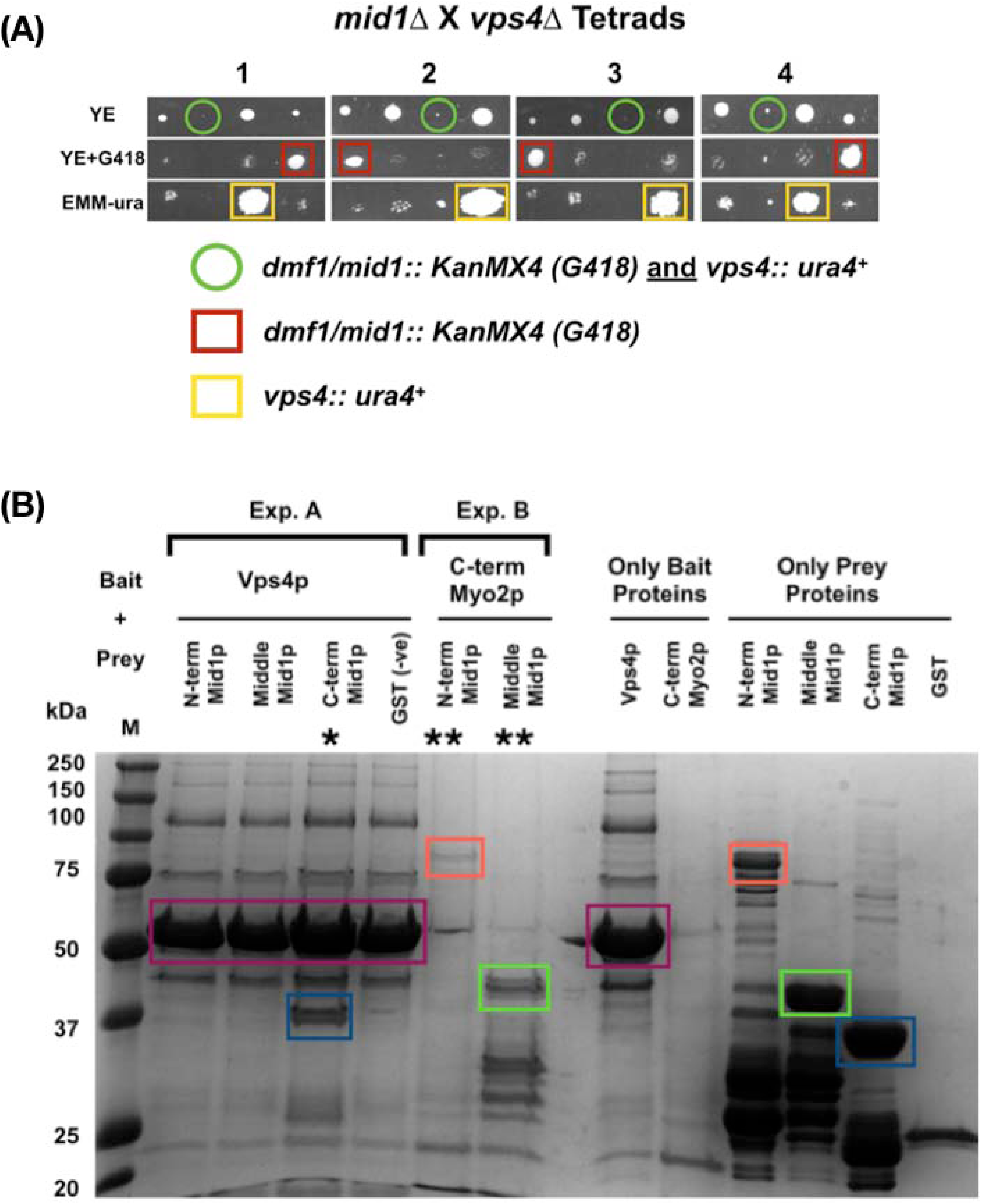
Genetic and physical interactions between *mid1* and *vps4* in fission yeast. **(A)** Synthetic lethality in *mid1*Δ *vps4*Δ double mutant cells indicates a genetic interaction between the *mid1* and *vps4* genes. Tetrad analysis of h*^-^ vps4*Δ (*vps4*::*ura4^+^*) mated with h*^+^ mid1*Δ (*dmf1*::*KanMX4*) to identify *vps4*Δ *mid1*Δ double mutants that show a synthetic lethal growth phenotype. Tetrads created by mating the two strains on solid ME medium, four spores dissected and allowed to grow on solid YE medium until colonies formed. Colonies replicated to solid YE+G418/KanMX4 and EMM-ura media and incubated to identify growth phenotypes and double mutants. **(B)** Direct physical interaction between Vps4p and the “C-term” domain of Mid1p. Recombinant GST-tagged (Mid1p domains: “N-term” (aa 1-453), “Middle” (452-579) and “C-term” (793-920)) and 6 His-tagged (Vps4p and “C-term” Myo2p) proteins were expressed in *E. coli* and purified using GST-sepharose beads and Ni-NTA agarose beads, respectively. Exp. A, eluted prey Mid1p domains were added to bait Vps4p bound to Ni-NTA agarose beads: (*) represents physical interaction of Vps4p and Mid1p “C-term” domain. Exp. B, eluted prey Mid1p “N-term” and “Middle” domains were added to bait Myo2p bound to Ni-NTA agarose beads as a positive control: (**) represents physical interaction of Mid1p “N-term” and “Middle” domains with “C-term” domain of Myo2p (left panel). The 14.5 kDa “C-term” Myo2p was not detected in this gel; in a separate experiment (right panel) the detected 14.5 kDa “C-term” Myo2p is shown. Asterisks (**) indicate detected physical interactions of Mid1p “N-term” and “Middle” domains with “C-term” domain of Myo2p. Coloured boxes represent predicted proteins: Vps4p purple, Mid1p “N-term” red, Plk1 purple, Mid1p “Middle” green and Mid1p “C-term” dark blue.

This striking observation demonstrated a genetic interaction between the *mid1* and *vps4* genes, with one explanation that the two encoded proteins interact to control an essential cellular function. To test this hypothesis we expressed and purified a His-tagged version of Vps4p protein from bacteria, using the same method to express and purify three different GST-tagged domains of Mid1p. Full length Mid1p was not purified as it is insoluble under such conditions [16, 18].

Full length Mid1p is 920 amino acids in length; the three domains used in pull-down experiments described here encompass the amino acids 1-453 (“N-term”), 452-579 (“Middle”) and 798-920 (“C-term”) [19, 18]. Of these three domains, only the C-terminus of amino acids 798-920 was pulled-down by recombinant Vps4p (Fig 1B). This indicates a direct physical interaction between Vps4p and the C-terminus of Mid1p, and that this interaction involved residues 798-920 of Mid1p.

### Vps4p is required for the correct cellular distribution of Mid1p

To further understand the interaction between Vps4p and Mid1p we examined the requirement for Vps4p to control the cellular distribution of GFP-tagged Mid1p during the cell cycle. Mid1p distribution has been well characterised and shown to move from the nucleus to the equatorial region to form nodes and a medial ring, that predicts the site of cell cleavage during septation [8].

Wild-type cells showed three patterns of GFP-Mid1p: localization to the nucleus, cytoplasmic, and equatorial nodes, as previously reported (Fig 2A; [8, 5]). By contrast, GFP-Mid1p in *vps4*Δ cells showed strikingly different patterns. Although cytoplasmic GFP-Mid1p was similar to wild-type, additional plasma membrane localization was observed (Fig 2B). In some cells, GFP-Mid1p localized to only one node and the plasma membrane, or localized to three nodes in other cells. The frequencies of these abnormal phenotypes in *vps4*Δ cells were compared with wild-type and showed significant differences (quantified in Fig 2C). Overall, GFP-Mid1p localization was significantly altered by the absence of *vps4^+^*.

**Fig 2.**
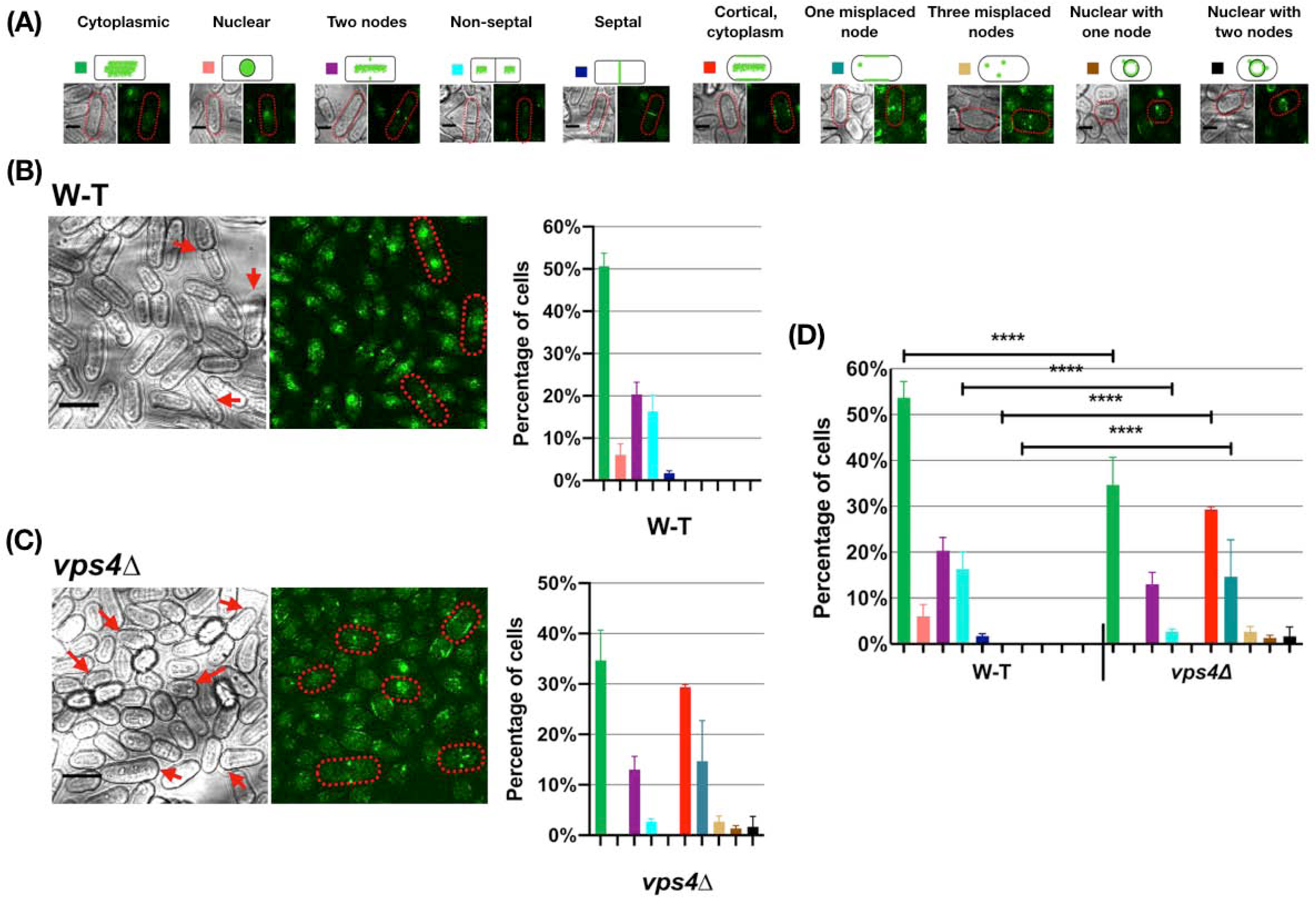
GFP-Mid1p cellular localization is disrupted in *vps4*Δ cells. Wild-type (W-T) and *vps4*Δ (*vps4*::*ura^+^*) *S. pombe* strains containing GFP-Mid1p grown at 25°C in liquid YE medium to mid-exponential phase and visualized by confocal microscopy. **(A)** Key to characterised GFP-Mid1p localization phenotypes. Scale bar, 5 µm. Bright and green fluorescent images of cells (left) and quantitative analysis (right) of GFP-Mid1p localization phenotypes for wild-type (W-T) **(B)** and *vps4*Δ **(C)** cells. Scale bar, 10 µm. **(D)** Two-way ANOVA analysis of frequencies of localization phenotypes in *vps4*Δ compared to wild-type (W-T). Asterisks (****) denote *p* values <0.0001 indicating significant differences to wild-type. Error bars, SEM.

These results, in addition to the observation that the *mid1* and *vps4* genes interact genetically, and further that the Mid1p and Vps4p physically interact *in vitro*, suggest that Mid1p and Vps4p coordinate to regulate the *S. pombe* cell cycle.

### Genetic and physical interactions between Mid1p and Ark1p in fission yeast

We hypothesized that *mid1* and *ark1* genes interact to accomplish *S. pombe* cytokinesis, as we and others have shown that aurora kinases modulate ESCRT protein function [20, 15]. To test this, both genetic and biochemical approaches were taken.

First, a *mid1*Δ strain was crossed with strains containing *ark1*-TS mutations to generate double *mid1*Δ *ark1*-TS mutant strains. The *mid1*Δ *ark1*-TS double mutant failed to form viable colonies (Fig 3A). Such a synthetic lethality indicates a genetic interaction between the *mid1* and *ark1* genes.

**Fig 3.**
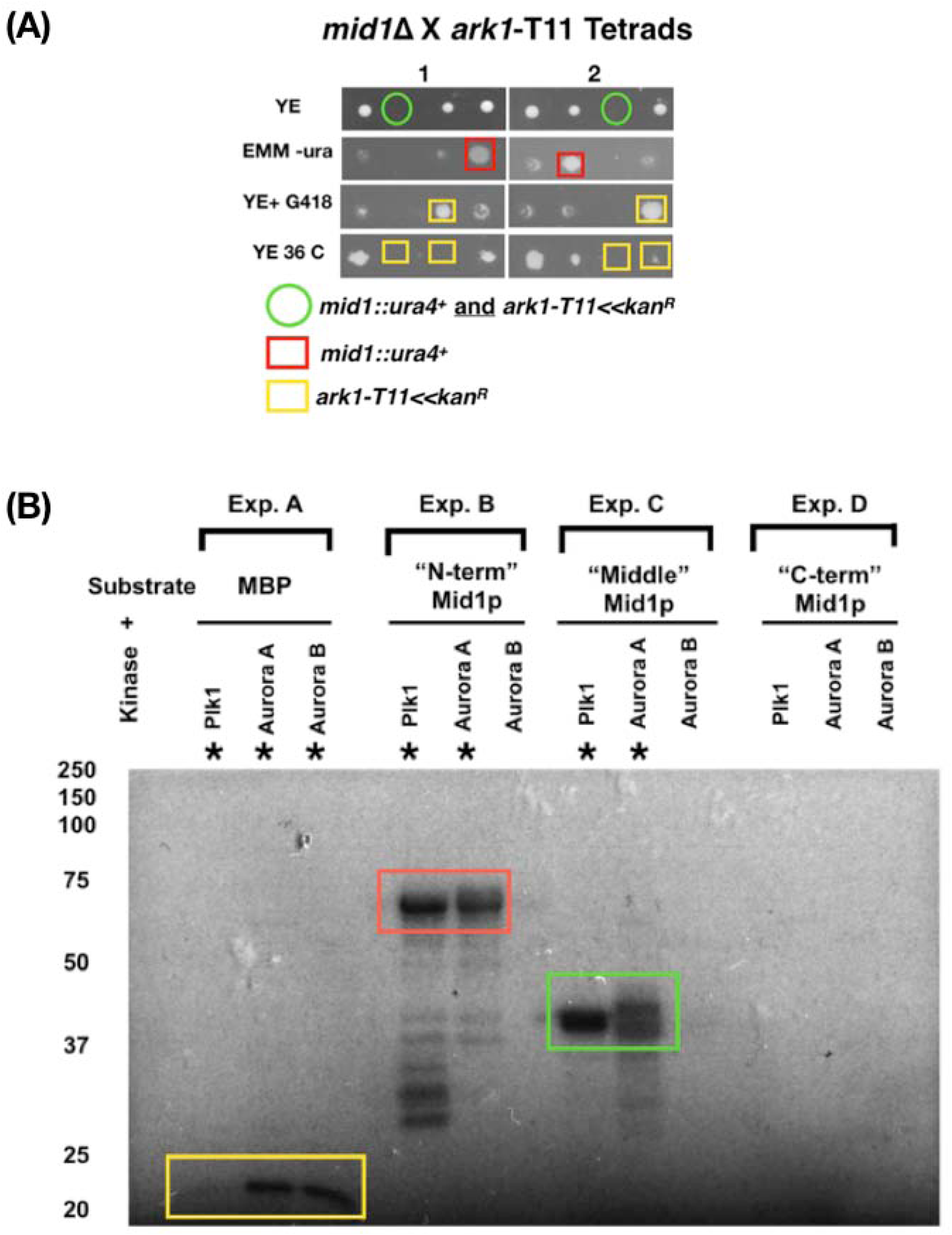
Genetic and physical interactions between *mid1* and *ark1* in fission yeast. **(A)** Synthetic lethality in *mid1*Δ *ark1*-T11 double mutant cells indicates genetic interaction between *mid1^+^* and *ark1^+^* genes. Tetrad analysis of h*^+^ ark1-T11* (*ark1-T11<<Kan^R^*) crossed with h*^-^ mid1*Δ (*mid1*::*ura^+^*) to identify *mid1*Δ *ark1-T11* double mutants that show a synthetic lethal growth phenotype. Tetrads created by mating the two strains on solid ME medium, four spores dissected and allowed to grow on solid YE medium until colonies formed. Colonies replicated to solid YE+G418/KanMX4 and EMM-ura media and incubated to identify growth phenotypes and double mutants. **(B)** Plk1 and Aurora A kinases phosphorylate Mid1p “N-term” and “Middle” domains. Recombinant GST-tagged Mid1p domain proteins “N-term” (aa 1-453), “Middle” (452-579) and “C-term” (798-920) were purified and analysed by *in vitro* phosphorylation assays. In “Exp. A” myelin basic protein (MBP) was a positive control. In “Exp. B”, “Exp. C” and “Exp. D” the “N-term”, “Middle” and “C-term” domains of Mid1p were substrates with the kinases Plk1, Aurora A or Aurora B, respectively. Asterisks (*) indicate detected *in vitro* phosphorylation signals. Coloured boxes represent predicted proteins: MBP yellow, Mid1p “N-term” species red, and Mid1p “Middle” species green.

As with *mid1* and *vps4* such a genetic interaction might be explained by a physical interaction between Mid1p and Ark1p proteins. Since Ark1p is a protein kinase we tested whether Mid1p can be phosphorylated by aurora kinase, using an *in vitro* phosphorylation approach with recombinant Mid1p domains purified from *E. coli*. These experiments were performed with the human aurora kinases Aurora A, Aurora B, and the human polo kinase Plk1 (Fig 3B). Phosphorylation of both the “N-term” and “Middle” Mid1p domains by Aurora A and Plk1 was detected, but not by Aurora B (Fig 3B; Exp. B and Exp. C). In contrast, no phosphorylation was detected of the “C-term” Mid1p domain by any of the three kinases Aurora A, Aurora B or Plk1 (Fig 3B; Exp. D). These results indicate that, at least *in vitro*, the “N-term” and “Middle” domains of Mid1p interact with and are phosphorylated by both Aurora A and Plk1 kinases, while the “C-term” domain of Mid1p is not phosphorylated by Aurora A, Aurora B or Plk1.

### Ark1p is required for the correct cellular distribution of Mid1p

To further analyse the interaction of Ark1p and Mid1p we examined the effect of two different temperature sensitive *ark1* mutants, *ark1-*T11 and *ark1-*T8, on Mid1p distribution in cells (Fig 4).

**Fig 4.**
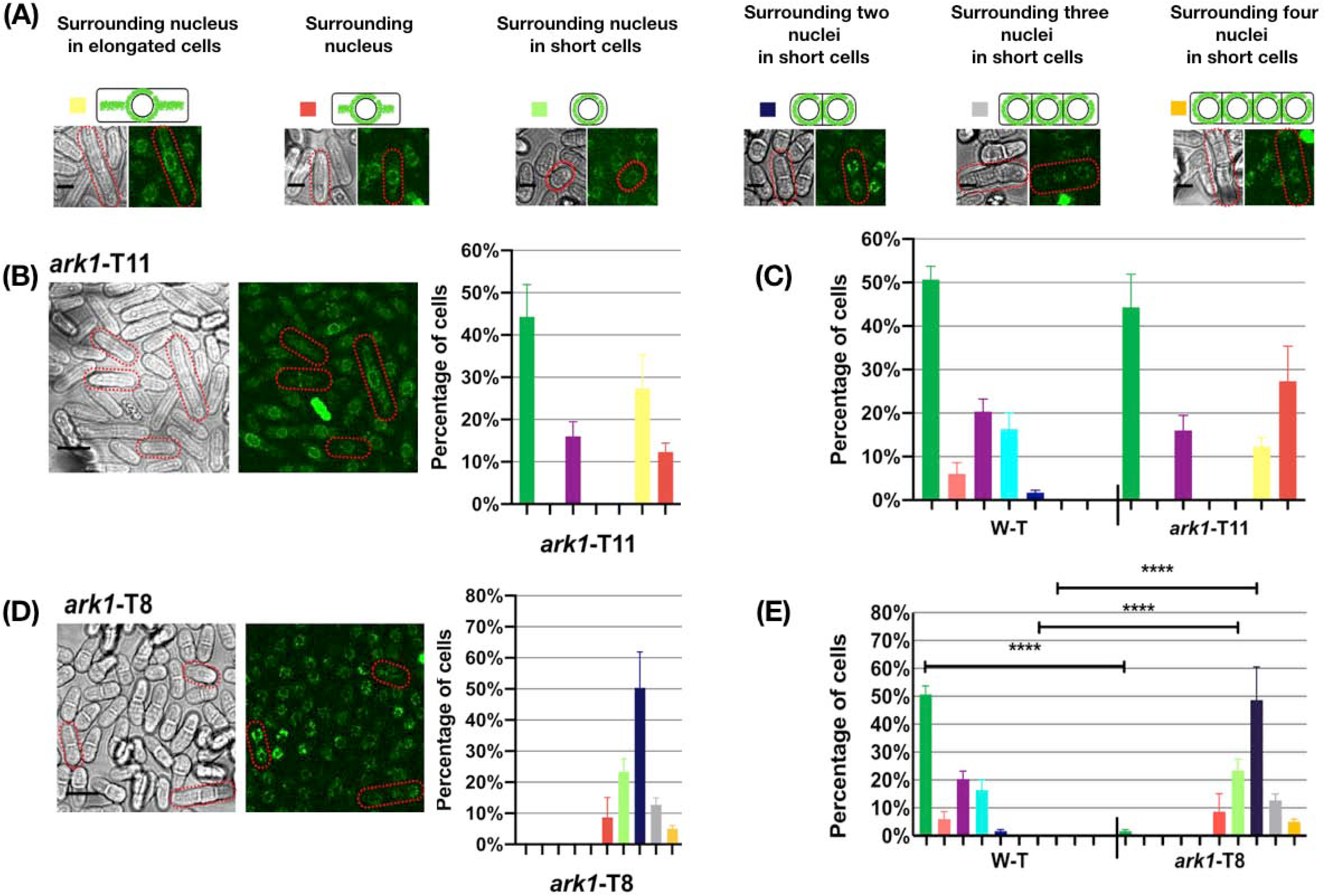
GFP-Mid1p cellular localization is disrupted in *ark1*-T11 and *ark1*-T8 cells. Wild-type (W-T), *ark1*-T11 and *ark1*-T8 *S. pombe* strains containing GFP-Mid1p were grown at 25°C in liquid YE medium to mid-exponential phase and visualized by confocal microscopy. **(A)** Key to characterised GFP-Mid1p localization phenotypes. Scale bar, 5 µm. Bright and green fluorescent images of cells (left) and quantitative analysis of GFP-Mid1p localisation phenotypes (right) for *ark1*-T11 **(B)** and *ark1*-T8 **(D)**. Scale bar, 10 µm. **(C, E)** Two-way ANOVA analysis of frequencies of localization phenotypes in *ark1*-T11 and *ark1*-T8 compared to wild-type (W-T). Asterisks (****) denote *p* values <0.0001 indicating a significant difference to wild-type. Error bars = SEM.

In *ark1*-T11 cells GFP-Mid1p showed wild-type phenotypes (Fig 4A). But in addition, new localization patterns were apparent, where GFP-Mid1p localization showed nuclear exclusion and appeared to be encircling the nucleus. The frequencies of these phenotypes were quantified and compared to wild-type cells revealing significant differences (Figs 4B). In *ark1*-T8 cells GFP-Mid1p showed new additional localization patterns (Fig 4D). Cells were observed to be round and shorter than the rod-shaped wild-type cells and GFP-Mid1p localization presented nuclear exclusion and encircled the nucleus. While some cells had one nucleus, others showed two, three, or even four nuclei. The frequencies of these phenotypes were quantified and compared to wild-type cells revealing significant differences (Figs 4E). These data support the hypothesis that Mid1p and Ark1p coordinate to regulate the *S. pombe* cell cycle.

### Identification of Mid1p amino acid residues phosphorylated by aurora and polo kinases

To further explore phosphorylation of Mid1p by aurora and polo kinases a mass spectrophotometry approach was used to identify phospho-acceptor sites. The “N-term” and “Middle” domains of Mid1p gel fragments from *in vitro* kinase assay reactions identical to those shown Fig 3B, but with non-labelled ATP, were excised and subjected to nano-scale liquid chromatographic tandem mass spectrometry (nLC-MS/MS) to generate a Mid1p phospho-site map (Fig 5). This analysis identified 35 potential Mid1p residues phosphorylated by either Plk1 or Aurora A kinases (S2 Table). In parallel we examined other published databases on fission yeast phospho-proteomes to refine and confirm the number of Mid1p phospho-sites. Most of these studies used a stable isotope labeling by amino acid in cell culture approach. We selected four studies which studied the *S. pombe* global proteome and identified several Mid1p specific phospho-sites [21–24].

**Fig 5.**
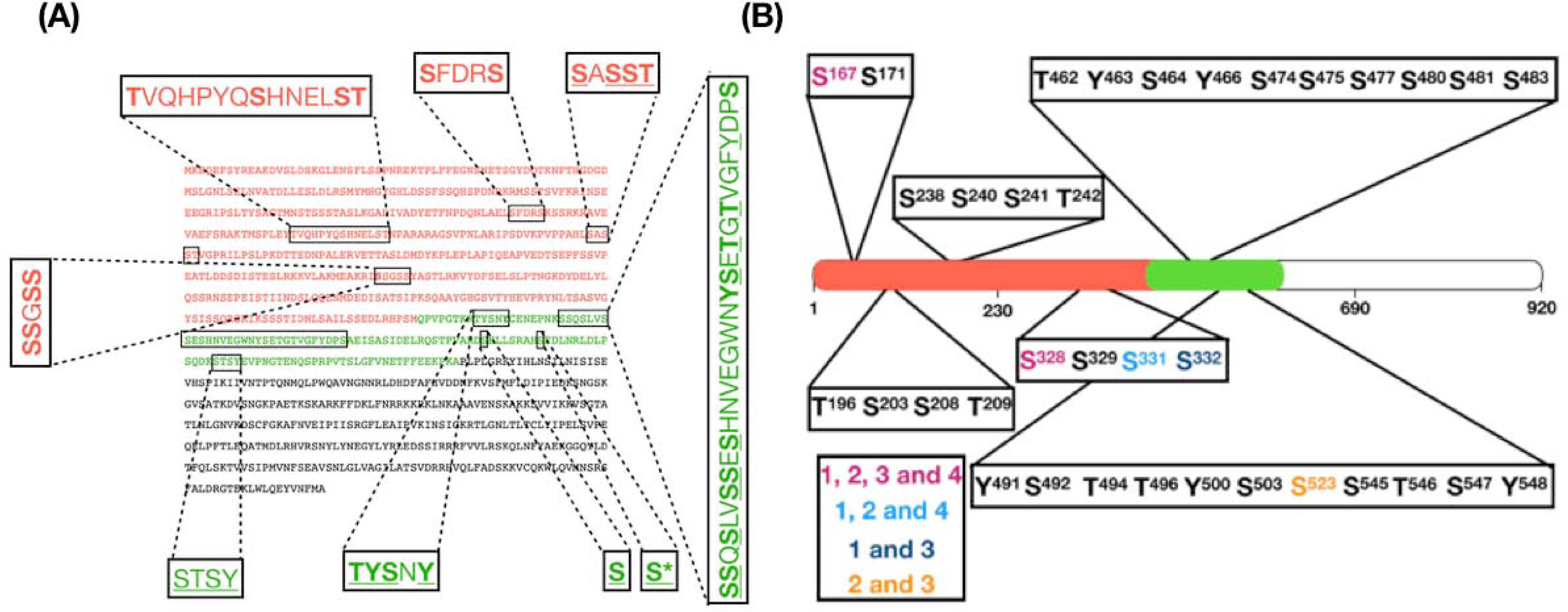
Identification of Mid1p phospho-sites by mass-spectrometry analysis (MSA) and published studies. **(A)** Mid1p full-length amino acid sequence with “N-term” and “Middle” domains indicated in red and green. Highlighted peptides show 35 phosphorylated amino acid residues identified by MSA of *in vitro* phosphorylation reactions for the kinases Plk1 (bold), Aurora A (underlined) or both (bold and underlined). **(B)** Five Mid1p phospho-sites generated from comparison of the 35 MSA amino acids with Mid1p phosphorylated amino acids identified in four published proteomic studies [21–24]. Amino acids identified by MSA and four independent studies (pink), MSA and three studies (light blue), and MSA and two studies (dark blue and orange). Asterisk (*) indicates a sixth serine phospho-site at position 531 derived from [23] and [24].

The Mid1p phosphorylation events identified by mass spectrophotometry are summarized in S3 Table, alongside those described by the four published studies. The red highlighted residues represent overlap with the residues identified reported here. The phospho-sites map shows a total of six potential phospho-sites distributed along the “N-term” and “Middle” Mid1p sequence, five of which were identified in this work, and the sixth was added from the published studies (Fig 5B). These are the serine residues S167, S328, S331, S332, S523 and S531.

To confirm the new phospho-sites identified in Mid1p, *in vitro* phosphorylation experiments involving bacterially expressed phospho-resistant mutant (serine to alanine) forms of the “N-term” and “Middle” domains of Mid1p with Aurora A and Plk1 kinases were completed (Fig 6). These experiments reveled markedly reduced phosphorylation of the Mid1p phospho-resistant mutant S167A by both Aurora A and Plk1 kinases, and S523A by Aurora A alone. These observations support the suggestion that the S167 and S523 residues are phosphorylated by the Aurora A and Plk1 kinases, at least *in vitro*.

**Fig 6.**
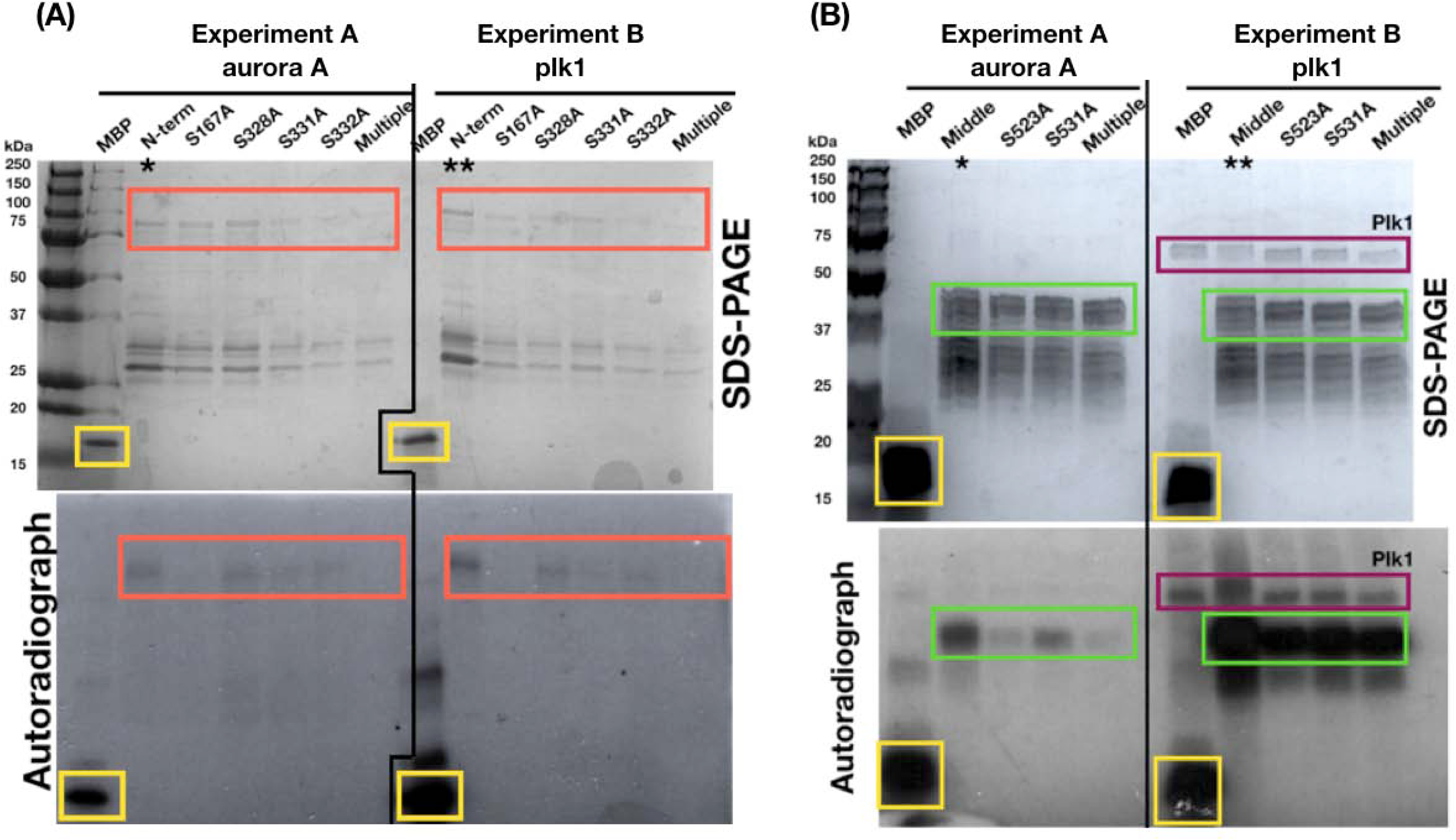
Reduced *in vitro* phosphorylation of Mid1p “N-term and “Middle” phospho-resistant mutants by Plk1 and Aurora A kinases. Recombinant GST-tagged Mid1p “N-term” **(A)** and “Middle” **(B)** domains phospho-resistant mutants were expressed and purified in *E. coli*. Proteins subjected to *in vitro* phosphorylation experiments with Aurora A (Exp. A) or Plk1 (Exp. B) kinases. Asterisks (* and **) indicate wild-type Mid1p “N-term” and Mid1p “Middle” recombinant proteins used as positive controls. Coloured boxes represent predicted proteins: MBP yellow, Mid1p “N-term” species red, Plk1 purple, and all Mid1p “Middle”-related species green. “Multiple” indicates mutations in all four phospho-sites.

### Biological relevance of amino acid residues phosphorylated by Ark1p and Plo1p in Mid1p

As another way to examine the role of the phospho-sites identified in Mid1p, we analysed the effect of their mutation *in vivo*. *S. pombe* cells exhibit morphology defects in the absence of Mid1p function. Therefore, we tested the effect of mutations of Mid1p phospho-sites in fission yeast on cell morphology to assay their relevance.

Versions of the *mid1* gene containing phospho-site mutations were generated and integrated in single copy into chromosomal DNA of *S. pombe mid1*Δ cells. Each version of the *mid1* gene had either a phospho-mimetic (S>D) or a phospho-resistant (S>A) point mutation(s) of the residues S167, S328, S331, S332, S523 or S531 to create a panel of mutant *S. pombe* strains (S1 Fig). Such *mid1* mutations were made both singly and in combination. As a positive control we integrated the wild-type *mid1*^+^ gene into *mid1*Δ cells to create *mid1*Δ pJK148:*mid1^+^*. This resulted in cells that behaved identically to wild-type both on solid medium and in liquid culture (Fig 7 and S1 Fig).

**Fig 7.**
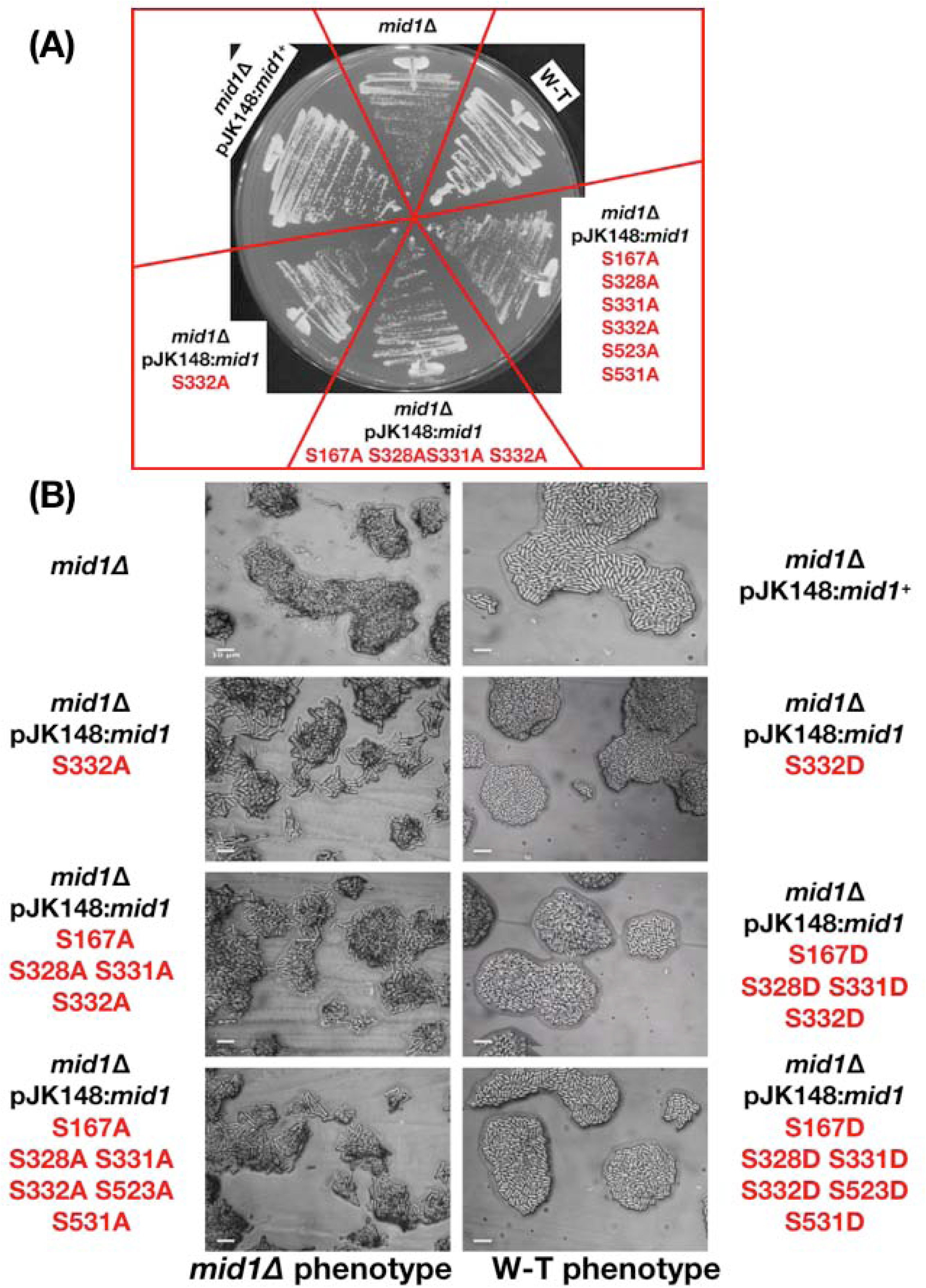
The S332 phospho-site is required for Mid1p function. Three phospho-resistant *mid1* mutant strains *mid1* S332A, *mid1* S167A S328A S331A S332A, and *mid1* S167A S328A S331A S332A S523A S531A, and the three phospho-mimetic mutants *mid1* mutant strains S332D, *mid1* S167D S328D S331D S332D, and *mid1* S167D S328D S331D S332D S523D S531D streaked to single colonies on solid YE at 25°C to compare growth rates **(A)** and colony morphology **(B)**. Controls: *mid1*Δ and wild-type (W-T) cells. Scale bar, 10 µm.

#### Mid1p phospho-mutants in mid1Δ cells

Initial experiments examined in *mid1*Δ cells the effect of Mid1p phospho-mimetic or phospho-resistant mutations. Strains were grown on solid medium, with the formation of colonies observed in all cases. In most strains where serine was changed to either alanine or aspartic acid cell morphology appeared similar to wild-type (Fig 7, and data not shown). However, interestingly, *mid1* S332A cells (but not *mid1* S332D) had defects in morphology similar to those observed in *mid1*Δ cells, with slower growth at 25°C (Fig 7). These data indicate a requirement of S332 for Mid1p function.

Furthermore, other phenotypes were revealed when certain Mid1p phospho-mutants were cultured in liquid medium. For example, *mid1* S523A and *mid1* S531A mutants displayed morphology defects (S2 Fig). In contrast, the equivalent phospho-mimetic mutants *mid1* S523D or *mid1* S531D did not.

#### Mid1p phospho-mutants in ark1-T11 cells

To examine the role of Mid1p phosphorylation in its interaction with Ark1p, we crossed the strains containing phospho-mimetic/resistant versions of *mid1* with *ark1*-T11 mutant cells and searched for synthetic phenotypes in double mutants. Viable colonies were observed in all cases with no synthetic phenotypes detected in most double mutants (data not shown). However, *mid1* S523A *ark1*-T11 and *mid1* S531A *ark1*-T11 double mutants showed cell morphology defects when cells were grown in liquid culture more severe than the single mutant strains (S3 Fig). These defects were not observed in the equivalent *mid1* S523D *ark1*-T11 or *mid1* S531D *ark1*-T11 double mutants. The cell morphology defects observed in *mid1* S523A *ark1*-T11 and *mid1* S531A *ark1*-T11 double mutants were defined as loss of the rod-like shape of cells. This suggests a role for these two phospho-sites in Mid1p function during the *S. pombe* cell cycle to ensure medial division plane placement, and consequently equal sized and rod-shaped daughter cells.

#### Mid1p phospho-mutants in plo1-ts35 cells

To examine the role of Mid1p phosphorylation in its interaction with Plo1p, we crossed strains containing phospho-mimetic/resistant versions of *mid1* with *plo1-ts35* mutant cells and searched for synthetic phenotypes in double mutants. Viable colonies were observed in all cases with no synthetic phenotypes detected for the majority of double mutants (data not shown). However, colony formation on solid medium was slower for *mid1* S332A *plo1-ts35* double mutants, and more severe cell morphology phenotypes were observed, compared to the single mutant strains (data not shown). These defects were not observed in the equivalent *mid1* S332D *plo1-ts35* double mutants. Such genetic interactions suggest a link between the regulation of Mid1p and Plo1p.

#### Mid1p phospho-mutants in vps4Δ cells

To examine the role of Mid1p phosphorylation in its interaction with Vps4p, we crossed strains containing phospho-mimetic/resistant versions of *mid1* with *vps4*Δ mutant cells and searched for synthetic phenotypes in double mutants, Viable colonies were observed in all cases with no synthetic phenotypes detected (data not shown). However, colony formation on solid medium was slower for *mid1* S523D S531D *vps4*Δ double mutants, and more severe cell morphology phenotypes were observed, compared to the single mutant strains. Strikingly, cells had defects in morphology similar to those observed in *mid1*Δ cells (Fig 8). The same serine residues changed to alanine S523A or S531A had no such effect when combined with *vps4*Δ (data not shown). These genetic interactions suggest a link between the regulation of Mid1p by both Vps4p and Ark1p.

**Fig 8.**
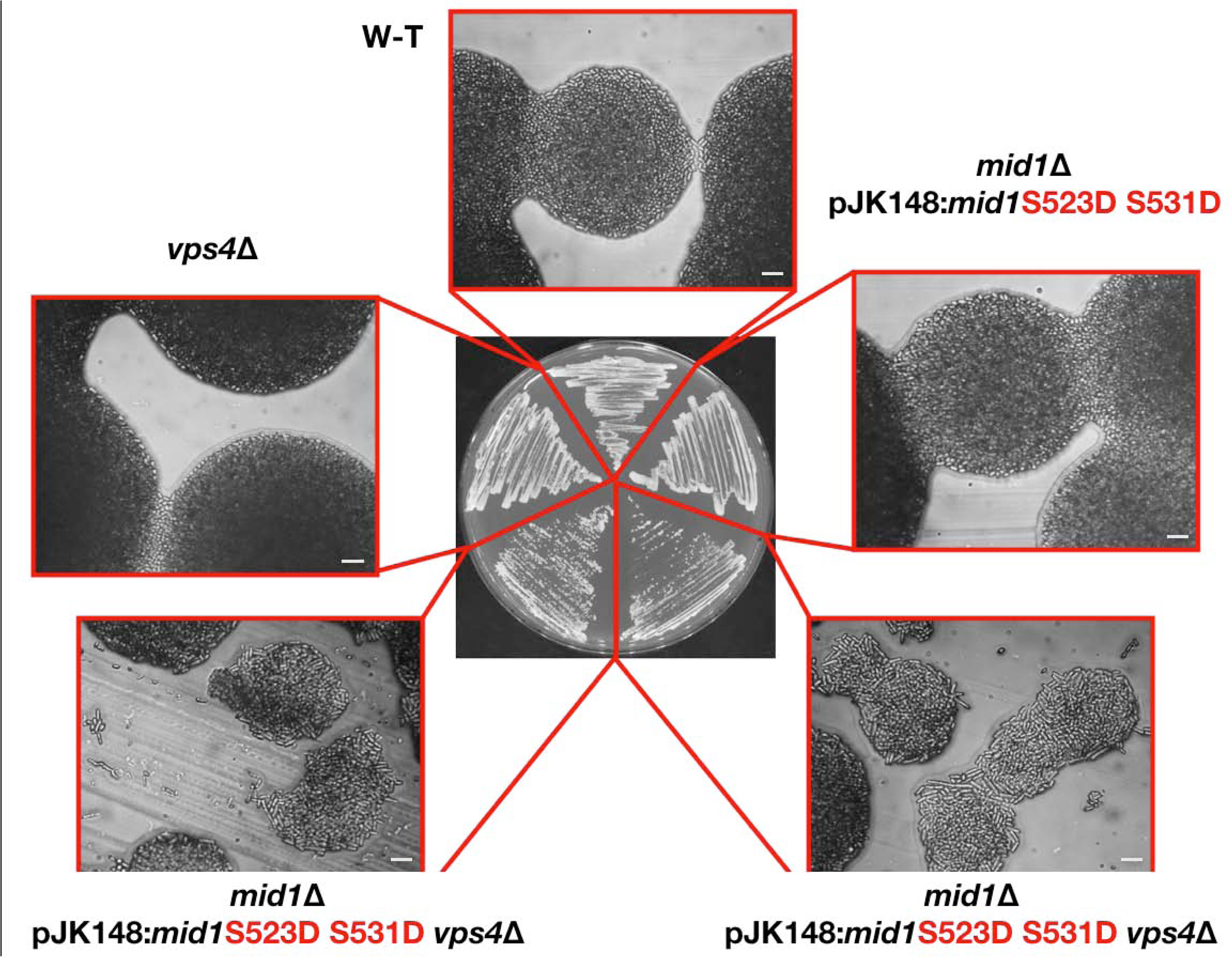
The Mid1p S523 and S531 phospho-sites regulate its interaction with Vps4p. Strains streaked to single colonies on solid YE medium and grown at 25° to compare growth rates and colony morphology. Controls: *vps4*Δ and wild-type (W-T) cells. Scale bar, 10 µm.

## Discussion

In fission yeast the Mid1p protein has important roles during cytokinesis during the cell cycle, with anillin homologues having similar roles in higher eukaryotes. These roles centre around its structural role in predicting and controlling the site of cell division in the equatorial region of the cell. Here we have identified new classes of proteins with which Mid1p interacts, and explored the mechanism by which these proteins cooperate and regulate each other and cell growth/morphology.

### Mid1p and Vps4p

Genetic and biochemical approaches revealed a direct physical interaction of Mid1p with the ESCRT-associated protein Vps4p. Furthermore, we demonstrated that a chromosomal deletion of the *vps4*^+^ gene caused defects in the cellular localization of GFP-Mid1p. These defects included mis-localization of nodes whereby one node is randomly positioned, or three nodes were present. This suggests a role of Vps4p in the Mid1p-dependent node localization pathway, which led us to hypothesize that the function of Mid1p is regulated by Vps4p during nodes attachment to the plasma membrane (Fig 9).

**Fig 9.**
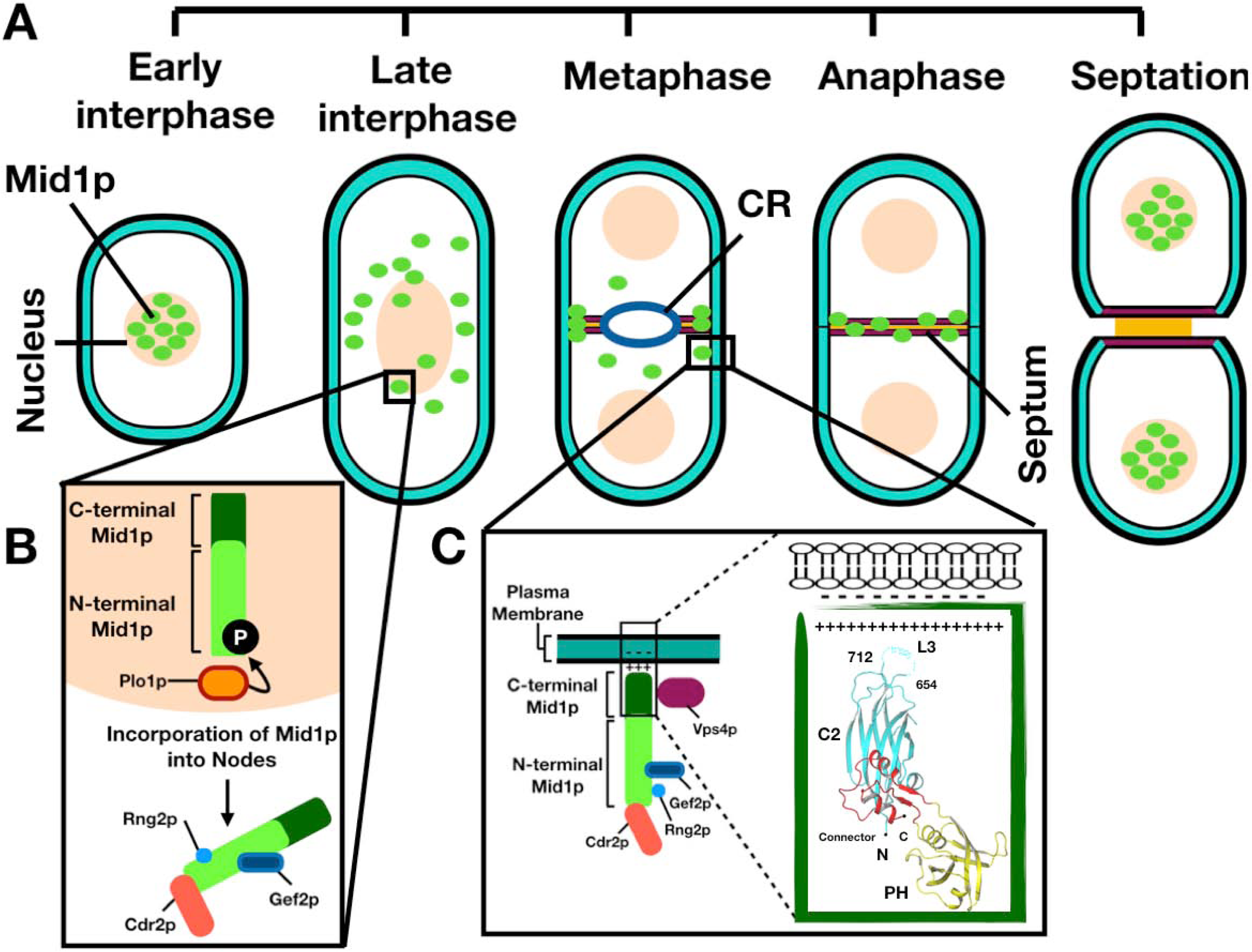
Interaction of Mid1p and Vps4p to regulate *S. pombe* septation. **(A)** Schematic representation of Mid1p localization during *S. pombe* cell cycle stages. CR = contractile ring. **(B)** Mid1p is phosphorylated by Plo1p to promote mitotic entry, during which Rng2p and other proteins interact with the N-terminal domain of Mid1p. **(C)** In mitosis, Mid1p forms nodes which are attached to the plasma membrane via the lipid binding motifs present within the Mid1p C-terminal domain. Green box represent the overall structure of Mid1p C-terminal region (aa 579-920) containing: the C2 domain (cyan), the connector domain (red) and the PH domain (yellow); dotted lines represent the lipid binding loop. Structure adapted from [13]. The Vps4p interaction with the C-terminal domain of Mid1p potentially regulates this process.

Two types of nodes are involved in *S. pombe* actomyosin ring assembly and contraction (Fig 9). Mid1p cortical anchorage first drives the recruitment of cytokinesis proteins, then interactions with myosin filaments causes the condensation of nodes into the actomyosin ring. Mid1p cortical anchorage depends on the PH domain [13] and its potential interaction with Vps4p might stabilize this interaction. Since Vps4p physically interacts with residues within the C-terminal domain of Mid1p, which contains membrane binding motifs, we speculate that Vps4p may facilitate Mid1p cortical anchorage to promote *S. pombe* medial division.

We suggest that a physical interaction between Vps4p and Mid1p regulates Mid1p-dependent node attachment to the plasma membrane to determine the division plane in *S. pombe*, and that this interaction directly or indirectly involves Mid1p PH domain cell cortex anchorage (Fig 9). It is interesting to note that the domain of Mid1p that interacts with Vps4p *in vitro* (Fig 1) contains the PH domain, suggesting that binding of Vps4p to this region may regulate interaction with the cell cortex (Fig 9).

### Phosphorylation of Mid1p

Genetic and biochemical approaches revealed a direct physical interaction of Mid1p with the aurora kinase, Ark1p, with Aurora A phosphorylating Mid1p the “N-term” and “Middle” domains. Mapping of the amino acids in Mid1p phosphorylated *in vitro* by Aurora A and Plk1 revealed 35 potential phospho-sites, with some of these sites independently identified in four *S. pombe* global proteomic studies (Fig 5). Such combined analysis suggested six potential phospho-sites in Mid1p: S167, S328, S331, S332, S523 and S531. Subsequent *in vitro* kinase assay experiments confirmed the *in vitro* phosphorylation of S167, S331 and S523 phospho-sites of Mid1p by Aurora A and Plk1 kinases (Fig 6).

To complement these studies, we examined the effect of mutations of these phospho-sites *in vivo* on cell morphology in wild-type, *ark1*-*T11, plo1-ts35* and *vps4*Δ S. *pombe* cells. In wild-type cells defective cell morphology phenotypes were observed for the phospho-mimetic mutants S332, S523 and S531 (Fig 7). These were exacerbated when combined with mutations in *ark1, plo1* and *vps4* (Fig 8). Therefore, we conclude that the phosphorylation of these amino acid residues is important for Mid1p function and its interaction with these proteins to regulate cell cycle events. It is tempting to speculate that the interaction of Mid1p and Vps4p is regulated by the activity of Ark1p and/or Plo1p, but it is important to note that the regions of Mid1p phosphorylated by these kinases do not include the binding region for Vps4p. Clearly, phosphorylation in adjacent regions may modulate binding via conformational changes in Mid1p, as regions containing the phospho-acceptor sites have been shown to regulate the interaction of Mid1p with other proteins, including Plo1p and Sid1p. Plo1p phosphorylates residues within the first 100 amino acids of Mid1p to trigger Myosin II recruitment during contractile ring assembly [25]. Later at contractile ring constriction Sid1p phosphorylates Mid1p to facilitate its export from the cell cortex [7]. Future experiments will be aimed at unraveling how the Mid1p interactome is modified both by association with Vps4p and by phosphorylation by mitotic kinases.

## Materials and Methods

### Yeast media and general techniques

General molecular procedures were performed, with standard methodology and media used for the manipulation of *S. pombe* [26, 27]. The yeast strains used in this study are shown in S1 Table. Cells were routinely cultured using liquid or solid complete (YE) or minimal (EMM) medium, at 25°C or 30°C.

### DNA constructs

The DNA constructs used in this study are listed in S2 Table. Some were synthesized and cloned by either GenScript or Invitrogen. All constructs were sequenced before use.

### Bacterial expression DNA constructs

Four fragments of the *mid1^+^* gene encoding the amino acids 1-453 “N-terminus”, 452-579 “Middle”, 578-799, 798-920 “C-term” [19] were synthesized and cloned into *Bam* HI/*Xho* I of pGEX-4T-1. Full-length *vps4^+^* was synthesized and cloned into *Nde* I/*Bam* HI of pET-14b. The C-terminal domain of *myo2^+^* was synthesized and cloned into *Nde* I/*Bam* HI of pET-14b.

### mid1 phospho-mimetic/resistant mutants

Eighteen versions of the *mid1* gene with different phospho-mimetic/resistant mutations were synthesized, along with a wild-type *mid1*^+^ control. All had the wild-type *mid1^+^* promoter in ∼1 kbp of DNA upstream of *mid1^+^* open reading frame, and were each cloned into *Kpn* I/*Sac* I of pJK148. Integration of the pJK148:*mid1* genes (1-19) into *S. pombe mid1*Δ cells was through homologous recombination after linearization of pJK148:*mid1* with *Nde* I in the *leu1*^+^ gene. The resulting panel of phospho-mimetic/resistant mutant strains is listed in S1 Table and S1 Fig.

### Recombinant protein purification

GST-Mid1p, 6His-Vps4p or 6His-C-term Myo2p plasmids were grown in BL21 *E. coli* until an OD of 0.6-0.8, with protein production induced by adding 1 mM IPTG. Mid1p 578-799 was not found to be expressed under these conditions, and so was not used for further experiments.

Bacterial pellets were produced by centrifugation with cells re-suspended in 20 ml re-suspension buffer with EDTA free protease inhibitors; for GST-Mid1p fusion proteins PBS buffer (137 mM NaCl, 2.7 mM KCl, 10 mM Na_2_HPO4 and 1.8 mM KH_2_PO4, pH 7.4) was used, whereas for 6His-Vps4p, HEPES buffer (25 mM HEPES, 400 mM KCl and 10% (v/v) glycerol, pH 7.4) was used. Cells were then lysed by sonication, where a final concentration of 1 mg ml^-1^ lysozyme was added for cell wall digestion followed by sonication, with 0.1 % (v/v) Triton X-100 added prior to lysis. A clear lysate was produced by centrifugation. GST-tagged fusion proteins were purified using 1ml l^-1^ glutathione beads in PBS buffer, while 6His-tagged Vps4p or C-term Myo2p were purified using 500 µl l^-1^ Ni-NTA beads in HEPES buffer, either for 2 h or overnight at 4°C. Mid1p was eluted from glutathione beads using Reduced glutathione buffer. Vps4p or C-term Myo2p were washed and eluted from Ni-NTA beads using HEPES buffers. Elution was carried out for 2 h at 4°C. Samples of proteins from each step were subjected to SDS-PAGE to determine elution efficiency.

### Pull-down experiments

Pull-down experiments utilized Ni-NTA beads-immobilized bait proteins (6His-Vps4p or 6His-C-term Myo2p) and prey-eluted proteins (GST-Mid1p: “N-terminus”, “Middle” or “C-terminus”) to investigate protein-protein interactions. Bait proteins xx ug were loaded onto Ni-NTA beads by incubation in PBS containing 0.01% (v/v) Triton X-100 for 1 h (4°C). After loading, the mixture was washed with PBS containing 0.01% (v/v) Triton X-100, and beads were blocked for non-specific binding by incubation in PBS containing 0.2% fish-skin gelatin. The beads mixture was incubated with yy ug prey protein in PBS with 0.01% (v/v) Triton X-100. Subsequently, beads were washed with PBS containing 0.01% (v/v) Triton X-100 three times, 0.5% (v/v) glycerol and 0.2% (w/v) fish skin gelatin three times, and with PBS alone four times. Proteins were eluted from beads by boiling in Laemmli sample buffer (LSB) (75 mM Tris pH 6.8, 12% (w/v) SDS, 60% (v/v) glycerol, 600 mM DTT and 0.6% (w/v) Bromophenol Blue) and samples were subjected to SDS-PAGE.

### In vitro kinase assays

Assays combined either human Plk1 (0.023 µg/µl), Aurora A (0.01 µg/µl), Aurora B (0.01 µg/µl) and myelin basic protein (MBP; 2.5 µg) (Sigma-Aldrich and Biaffin GmbH) or one of the three Mid1p domains (“N-term”, “Middle” or “C-term” 2.5 µg). Kinase, substrate proteins, 1 µCi [γ-^32^P] ATP, 10 mM ATP and kinase assay buffer (25 mM MOPS, 25 mM MgCl_2_, 1 mM EDTA and 0.25 mM DTT, pH 7.2) were mixed in a total volume of 20 µl; the reaction was initiated by adding 5 µl ATP cocktail and incubated at 30°C for 1 h and terminated by the addition of LSB; samples were subjected to SDS-PAGE followed by autoradiography or phospho-imaging. Following detection of *in vitro* phosphorylation signals, mass spectrometric analysis was carried out by the Dundee FingerPrints Proteomics service (http://proteomics.lifesci.dundee.ac.uk/).

### Confocal and light microscopy

Septation and GFP studies were carried out by visualizing *S. pombe* using calcofluor-white stain and fluorescence microscopy, respectively. *S. pombe* cells were cultured in 50 ml YE shaking at 28°C (25°C for *ark1*-TS strains) for 18 hours. Cell septa were visualized using bright field and DAPI filters of a Zeiss Axiovert 135 fluorescent microscope equipped with a Zeiss 63X Plan-APOCHROMAT oil-immersion objective lens. GFP-tagged *mid1^+^* [28] was examined by a He/Ne and Ag laser system of Zeiss LSM microscope using 63X high NA objective lens. Cell images were collected using Zeiss Pascal software and processed using ImageJ, Microsoft PowerPoint and Keynote software. Numerical analysis was completed from three independent experiments where 200-250 cells were counted. Yeast colonies grown on solid medium were imaged using a Zeiss Axioscope microscope and a Sony DSC-75 camera.

## Acknowledgements

We thank Anne Paoletti, Tomoko Iwaki and Silke Hauf for sharing fission yeast strains used in this study. We thank Ian Salt for his contribution to this project, both with experimental and practical advice. We acknowledge the following for contributions to experiments: Rianne Cort, Laura Downie, Bethany Hutton, Christina Jack, Jack Goddard, Susan McGill, Mark McLean, Chiamaka Okoli, Elena Pescuma, Liam Pollock, James Provan, John Riddell, Aizhan Shagadatova, Ellen Shercliff, Haoying Wang, and Wandiahyel Yaduma.

---

**S1 Fig.**
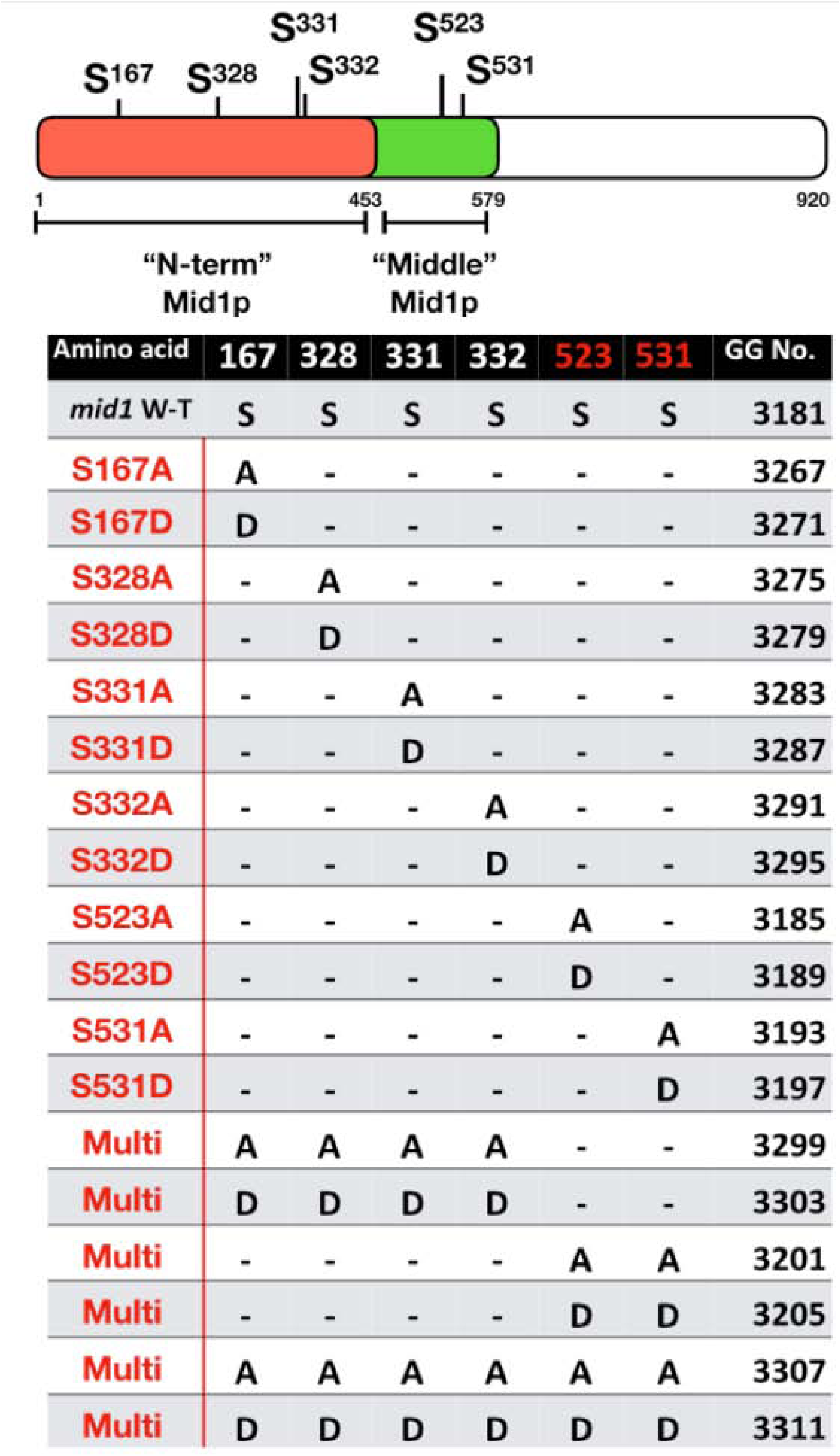
Summary of Mid1p phospho-mimetic/resistant fission yeast mutant strains used in this study. Top Panel: full length Mid1p with S167, S328, S331, S332, S523 and S531 phospho-sites indicated. Lower Panel: summary of single or combinatorial (“Multi”) phospho-mimetic or phospho-resistant mutations of the *mid1* gene, to generate eighteen versions integrated into *mid1*Δ cells, with each yeast strain given a “GG” lab collection number.

**S2 Fig.**
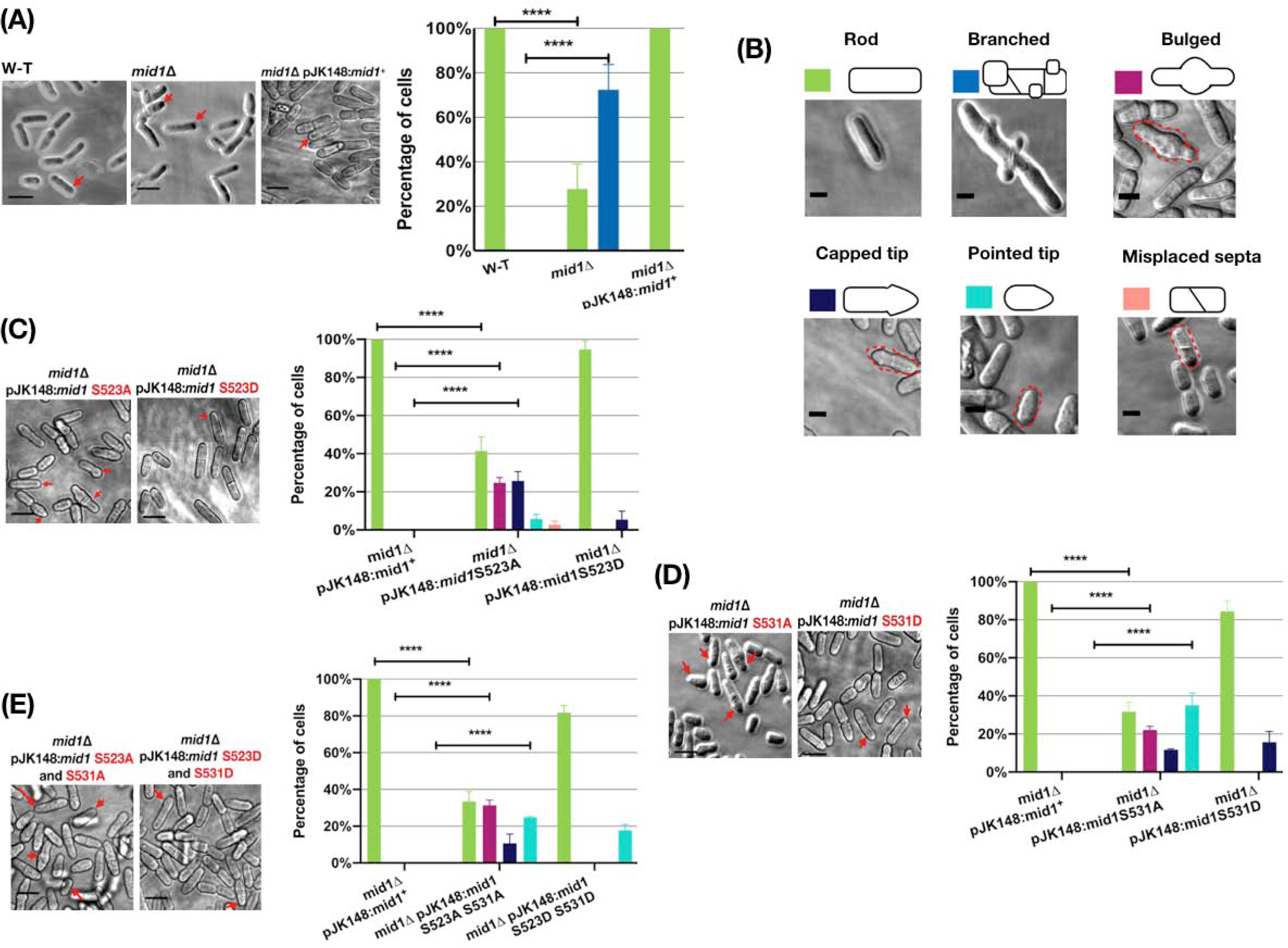
*mid1* phospho-resistant/mimetic mutants reveal cell division phenotypes. Wild-type (W-T), *mid1Δ*, *mid1Δ* pJK148:*mid1*^+^, *mid1Δ* pJK148:*mid1*S523A, *mid1Δ* pJK148:*mid1*S523D, *mid1Δ* pJK148:*mid1*S531A, *mid1Δ* pJK148:*mid1*S531D, *mid1Δ* pJK148:*mid1*S523A S531A and *mid1Δ* pJK148:*mid1*S523D S531D strains grown at 25°C in liquid YE medium to mid-exponential phase. Cells visualized by confocal microscopy under bright field. Scale bar, 10 µm. Characterised cell morphology phenotypes indicated by red arrows. **(A)** Two-way ANOVA analysis of frequencies of localization phenotypes in wild-type (W-T), *mid1Δ* and *mid1Δ* pJK148:*mid1*^+^ cells. **(B)** Key to characterised cell morphology phenotypes. Scale bar, 5 µm. **(C-E)** Two-way ANOVA analysis of frequencies of localization phenotypes of *mid1* phospho-resistant/mimetic mutants. Asterisks (****) denote *p* values <0.0001 indicating significant difference to wild-type. Error bars = SEM.

**S3 Fig.**
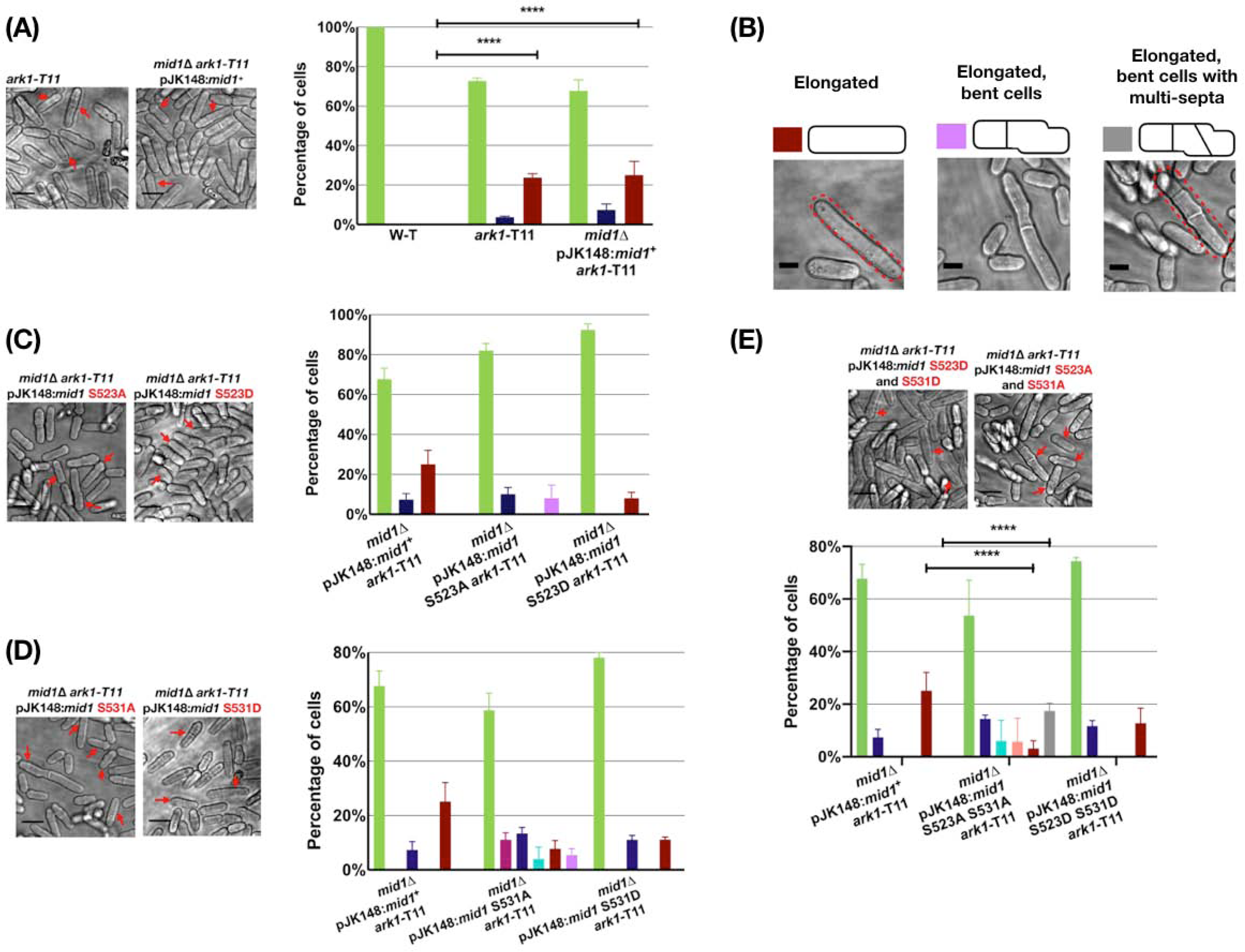
*mid1* phospho-resistant/mimetic *ark1*-T11 double mutants reveal cell division phenotypes. *ark1*-T11, *mid1Δ ark1*-T11 pJK148:*mid1*^+^, *mid1Δ ark1*-T11 pJK148:*mid1*S523A, *mid1Δ ark1*-T11 pJK148:*mid1*S523D, *mid1Δ ark1*-T11 pJK148:*mid1*S531A, *mid1Δ ark1*-T11 pJK148:*mid1*S531D, *mid1Δ ark1*-T11 pJK148:*mid1*S523A S531A and *mid1Δ ark1*-T11 pJK148:*mid1*S523D S531D strains grown at 25°C in liquid YE medium to mid-exponential phase. Cells visualized by confocal microscopy under bright field. Scale bar, 10 µm. Characterised cell morphology phenotypes indicated by red arrows. **(A)** Two-way ANOVA analysis of frequencies of localization phenotypes in wild-type (W-T), *ark1*-T11 and *mid1Δ* pJK148:*mid1*^+^ *ark1*-T11 cells. (**B)** Key to characterised cell morphology phenotypes. Scale bar, 5 µm. **(C-E)** Two-way ANOVA analysis of frequencies of localization phenotypes in wild-type (W-T), *mid1* phospho-resistant/mimetic and *ark1*-T11 cells. Asterisks (****) denote *p* values <0.0001 indicating significant difference to wild-type. Error bars = SEM.

**S1 Table.**
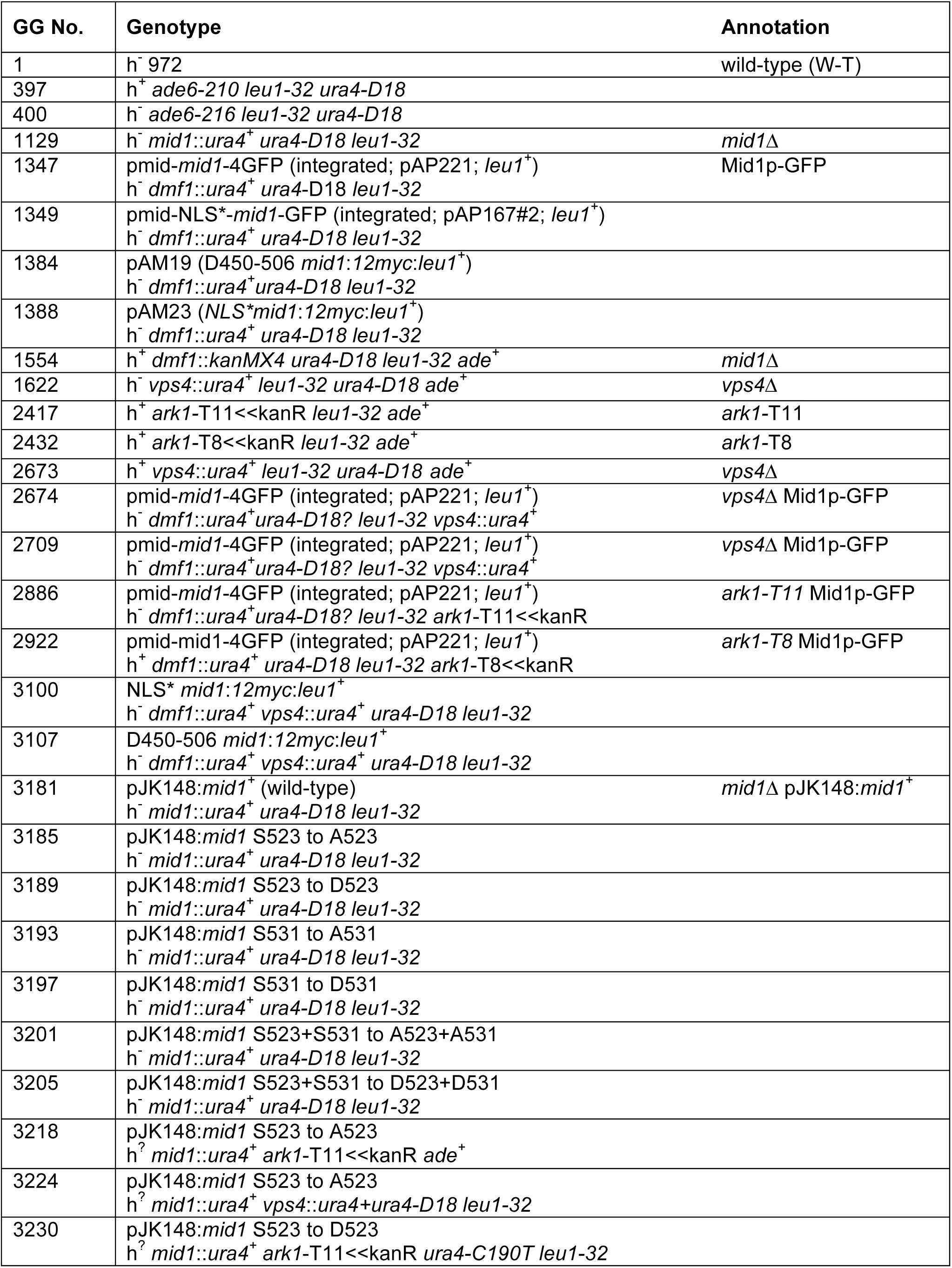
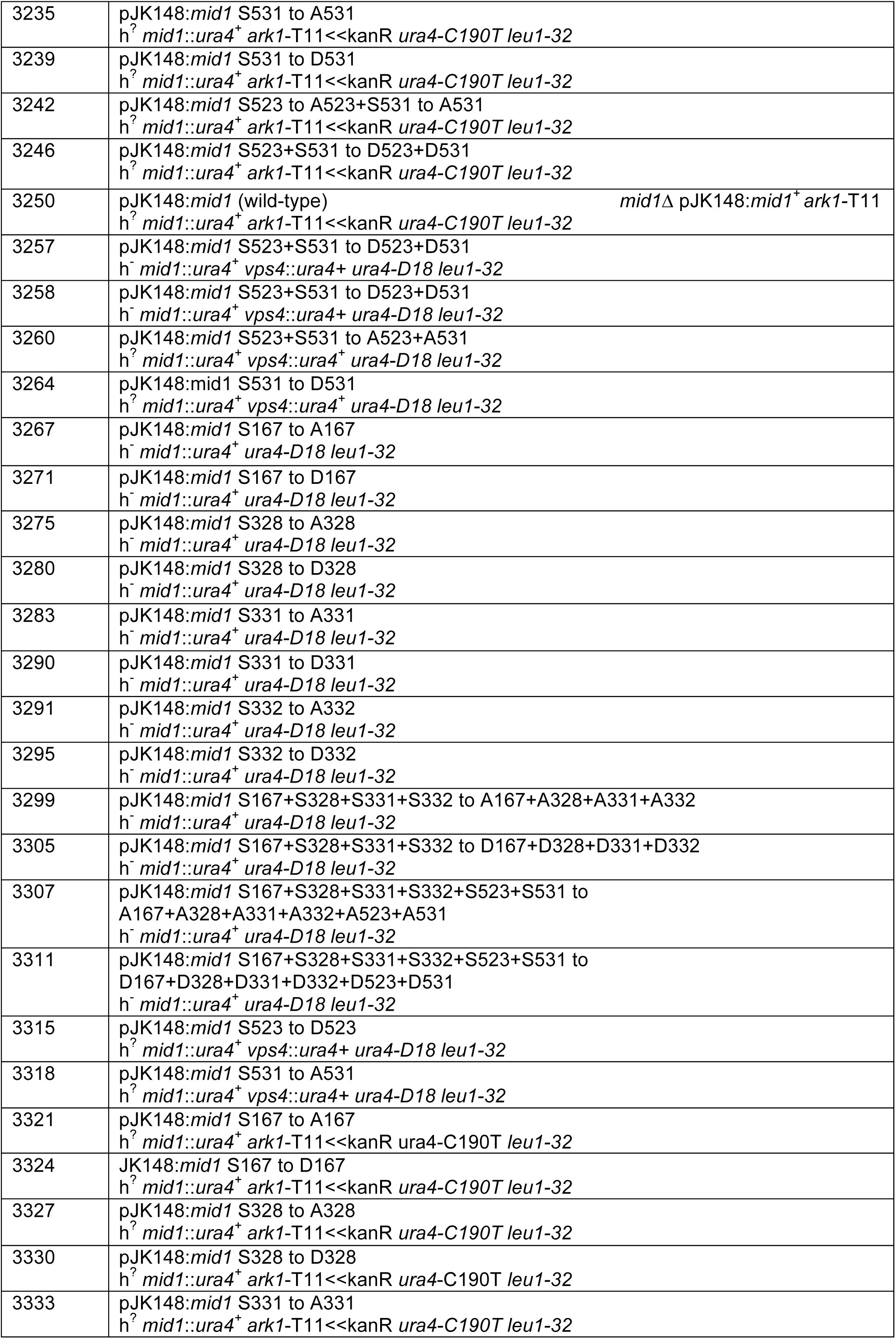
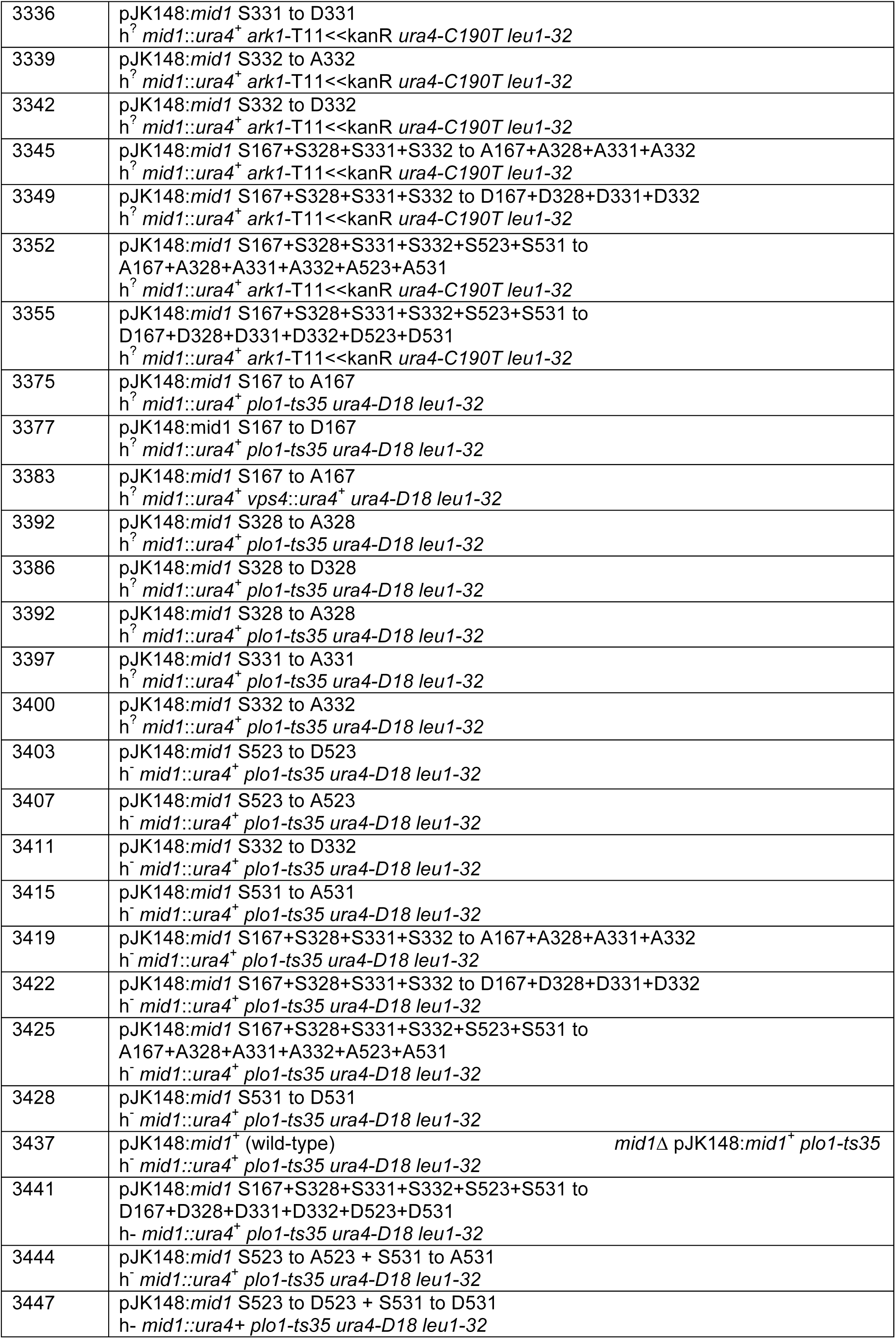
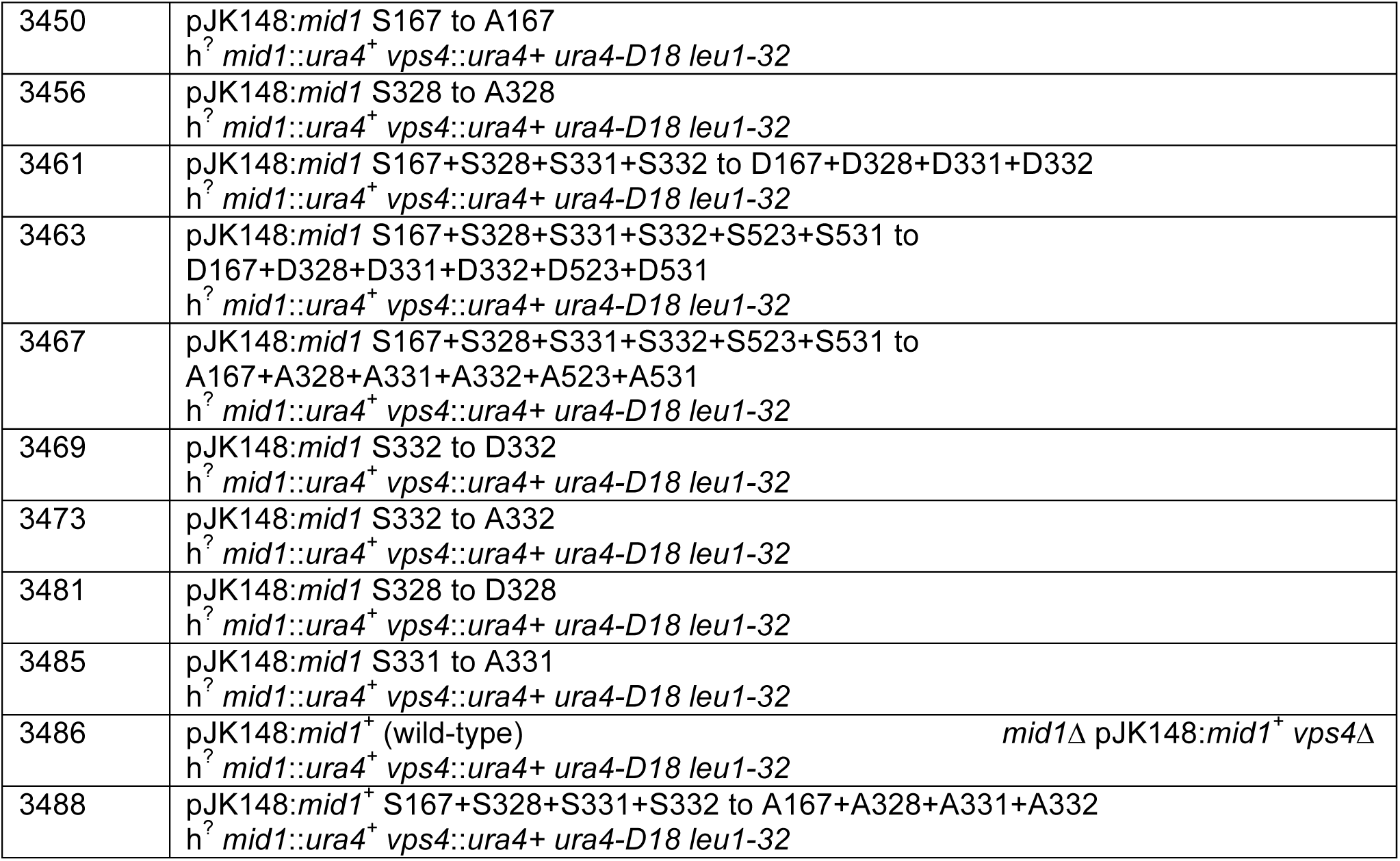
*S. pombe* strains used in this study. “GG” number refers to the laboratory reference collection. All strains *ade*^-^, unless indicated.

**S2 Table.** Vector DNA constructs used in this study. “GB” number refers to the laboratory reference collection.

| GB No. | Abbreviated plasmid vector description |
| --- | --- |
| 880 | pET-14b - His tagged, Vps4p full length, *Nde* I/*Bam* HI<br>Invitrogen order 13AB6ZFP |
| 881 | pGEX4T1 - GST tagged, Mid1p 1-453, *Bam* HI/*Xho* I - "N-term"<br>GenScript order U2640BJ110 |
| 882 | pGEX4T1 - GST tagged, Mid1p 452-579, *Bam* HI/*Xho* I - "Middle" |
| 883 | pGEX4T1 - GST tagged, Mid1p 578-799, *Bam* HI/*Xho* I |
| 884 | pGEX4T1 - GST tagged, Mid1p 798-920, *Bam* HI/*Xho* I - "C-term" |
| 889 | pET-14b - His tagged, Myo2p C-terminus amino acids 1394-1526, *Nde* I/*Bam* HI<br>GenScript order U9540CD270_2 |
| 907 | *mid1*<sup>+</sup> wild-type + 1 kb upstream of ORF - 3,853 base pairs in total<br>*Kpn* I/*Sac* I fragment cloned into pJK148<br>GenScript order U2002DH100 |
| 909 | *mid1* mutant (1) S523 to A523 in pJK148 |
| 911 | *mid1* mutant (1) S523 to D523 in pJK148 |
| 913 | *mid1* mutant (1) S531 to A531 in pJK148 |
| 915 | *mid1* mutant (1) S531 to D531 in pJK148 |
| 917 | *mid1* mutant (2) S523 to A523 + S531 to A531 in pJK148 |
| 919 | *mid1* mutant (2) S523 to D523 + S531 to D531 in pJK148 |
| 923 | *mid1* mutant (1) S167 to A167 in pJK148<br>GenScript order U894VEB050 |
| 924 | *mid1* mutant (1) S167 to D167 in pJK148 |
| 925 | *mid1* mutant (1) S328 to A328 in pJK148 |
| 926 | *mid1* mutant (1) S328 to D328 in pJK148 |
| 927 | *mid1* mutant (1) S331 to A331 in pJK148 |
| 928 | *mid1* mutant (1) S331 to D331 in pJK148 |
| 929 | *mid1* mutant (1) S332 to A332 in pJK148 |
| 930 | *mid1* mutant (1) S332 to D332 in pJK148 |
| 931 | *mid1* mutant (4) S167 to A167 + S328 to A328 + S331 to A331+ S332 to A332 in pJK148 |
| 932 | *mid1* mutant (4) S167 to D167 + S328 to D328 + S331 to D331+ S332 to D332 in pJK148 |
| 933 | *mid1* mutant (6) S167 to A167 + S328 to A328 + S331 to A331+ S332 to A332 + S523 to A523 + S531 to A531 in pJK148 |
| 934 | *mid1* mutant (6) S167 to D167 + S328 to D328 + S331 to D331+ S332 to D332 + S523 to D523 + S531 to D531 in pJK148 |
| 935 | pGEX4T1 - GST tagged, Mid1p 1-453, *Bam* HI/*Xho* I - "N-term" S167 to A167<br>GenScript order U8198EF120 |
| 936 | pGEX4T1 - GST tagged, Mid1p 1-453, *Bam* HI/*Xho* I - "N-term" S328 to A328 |
| 937 | pGEX4T1 - GST tagged, Mid1p 1-453, *Bam* HI/*Xho* I - "N-term" S331 to A331 |
| 938 | pGEX4T1 - GST tagged, Mid1p 1-453, *Bam* HI/*Xho* I - "N-term" S332 to A332 |
| 939 | pGEX4T1 - GST tagged, Mid1p 1-453, *Bam* HI/*Xho* I - "N-term"<br>S167 to A167 + S328 to A328 + S331 to A331 + S332 to A332 |
| 940 | pGEX4T1 - GST tagged, Mid1p 452-579, *Bam* HI/*Xho* I - "Middle" S523 to A523 |
| 941 | pGEX4T1 - GST tagged, Mid1p 452-579, *Bam* HI/*Xho* I - "Middle" S531 to A531 |
| 942 | pGEX4T1 - GST tagged, Mid1p 452-579, *Bam* HI/*Xho* I - "Middle"<br>S523 to A523 + S531 to A531 |

**S3 Table.**
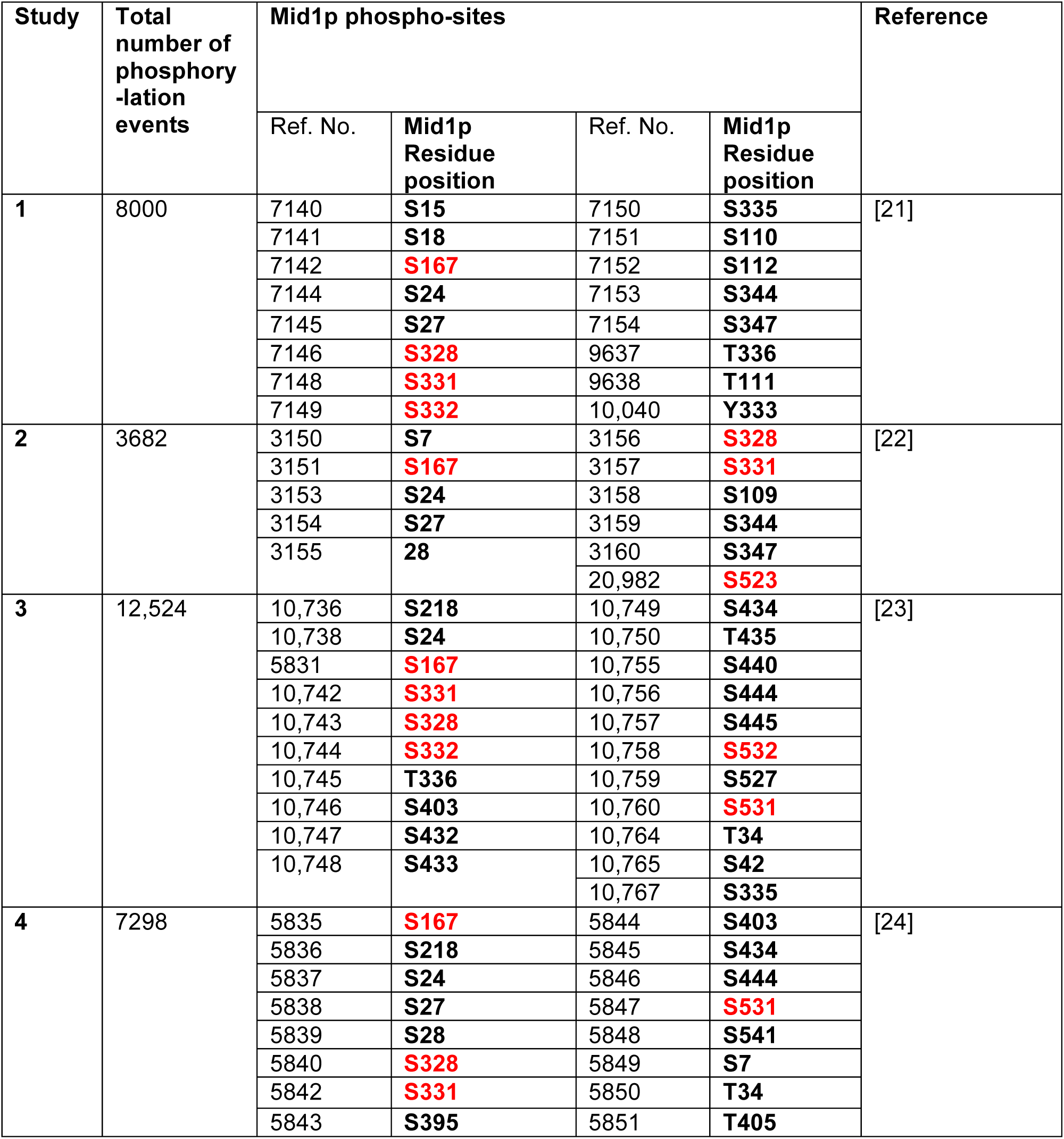
Summary of *S. pombe* global proteomic studies of identified Mid1p phospho-sites.

